# High-Throughput Screen of Advanced Repurposing Library Identifies Compounds Active Against Acute and Chronic Stages of *Toxoplasma gondii*

**DOI:** 10.64898/2026.11.13.766473

**Authors:** Taher Uddin, Case W. McNamara, Arnab K. Chatterjee, L. David Sibley

**Affiliations:** Department of Molecular Microbiology, Washington University School of Medicine, St. Louis, Missouri 63110, United States; Calibr-Skaggs Institute for Innovative Medicines, La Jolla, California 92037, United States

**Keywords:** High-Throughput Screening, Bradyzoites, ReFRAME

## Abstract

*Toxoplasma gondii*, a pathogen with a global burden impacting millions, has the capability to interconvert between the actively replicating tachyzoite form into a semi- dormant bradyzoite for, complicating treatment. Here, we utilized an advanced high- throughput screen (HTS) to identify repurposing candidates from the ReFRAME library, which consolidates 14,095 molecules from three major databases, including FDA- approved and late-stage clinical leads. To facilitate screening, the permissive cyst- forming *Tg*68 strain was genetically engineered to express nano luciferase under the pBAG1 promoter to track bradyzoite growth, and firefly luciferase under the pTub1 promoter to monitor tachyzoite growth. HTS in 384-well format identified 160 compounds that exhibited > 80% inhibition at 10 µM of both tachyzoites and bradyzoites. By cross-referencing against a publicly available HEK293T cytotoxicity dataset, we filtered out 61 compounds based on half-maximal cytotoxicity (CC_50_) less < 5 µM, leaving 99 for further evaluation. From this set, 15 top hits were identified based on potency and the selective index (CC_50_:EC_50_; SI > 20) against HepG2. The top hits were profiled using *in vitro* growth assays for tachyzoites, bradyzoites, and *ex vivo* liberated and intact cysts. Marizomib, monensin, narasin, and LY-2090314 showed the strongest inhibition following 24 h treatment of intact cysts. When tested in combination with clinical compounds atovaquone and pyrimethamine, the top hits were predominantly additive. Additionally, they were evaluated for their potential for developing resistance using a minimum inoculum of resistance (MIR) assay. Six of nine compounds tested in MIR assays yielded no detectable parasites over 30 days. In contrast, LY-2090314-resistant populations exhibited 26–29-fold resistance and carried a TgGSK3 L53V mutation, identifying a candidate resistance determinant. Collectively, our findings identify compounds with activity against tachyzoites and bradyzoites and with low propensity for resistance, as starting points for mechanistic investigation and further evaluation of *in vivo* efficacy and host-toxicity liabilities.

## Introduction

*Toxoplasma gondii* is an obligate intracellular parasite that establishes persistent infection through differentiation of rapidly replicating tachyzoites into bradyzoites. The semi-dormant bradyzoites are enclosed within tissue cysts, which preferentially persist in the brain, eyes, skeletal muscle, and cardiac muscle.^1–3^ Although chronic infection is often subclinical in immunocompetent individuals, reactivation in immunocompromised patients can cause severe disease.^4, 5^ Available therapies primarily control tachyzoite replication but do not eliminate tissue cysts, leaving infected individuals vulnerable to subsequent reactivation.^6, 7^ Treatment-associated toxicity and the need for prolonged therapy further emphasize the need for drugs that combine activity against acute and persistent parasite stages with acceptable host-cell selectivity.^6–8^.

Discovery of new active compounds against chronic infection is complicated by the distinct physiology of bradyzoites and the limitations of conventional tachyzoite-based screening.^9–12^ Bradyzoites are semi-dormant and their metabolic and replicative states can vary within and between tissue cysts, potentially influencing drug susceptibility.^13, 14^ Consequently, establishing that compounds are active against tachyzoites does not establish activity against persistent bradyzoite parasites.^3, 9, 15^ Screening approaches that incorporate differentiated bradyzoites and assess recovery after compound withdrawal are, therefore, important for identifying compounds that are effective against chronic infection.^9, 11, 16, 17^

Recent work from our group established a screening platform using the differentiation-permissive *T. gondii* isolate Tg68 cultured under alkaline stress or glucose withdrawal supplemented with glutamine to induce bradyzoite development *in vitro*.^9^ These conditions enable evaluation of compound activity in distinct differentiation- associated environments and provide an opportunity to identify susceptibility that would be missed by tachyzoite-only screening ^9^. Additionally, *ex vivo* evaluation of compound activity against bradyzoites recovered from tissue cysts provides an important benchmark for potential activity *in vivo* ^18–22^.

Drug repurposing offers a potential strategy for identifying new compounds that are active against *T. gondii* ^9, 17^. The ReFRAME (https://reframedb.org/) collection brings together approved drugs and investigational compounds with prior development histories, providing access to chemically and pharmacologically diverse molecules that can serve as repurposing candidates or mechanistic probes. Its utility for infectious disease and other discovery has been demonstrated by screens identifying compounds active against *Plasmodium*, *Cryptosporidium*, intracellular *Trypanosoma cruzi*, and SARS-CoV-2, highlighting its applicability across protozoan and viral pathogens.^23–28^ Here, we screened 14,095 ReFRAME compounds against tachyzoites and in vitro bradyzoites induced under two conditions using the differentiation-permissive Tg68. We further identified 15 top hits through secondary profiling, *ex vivo* assays of liberated bradyzoites and intact tissue cysts, drug-combination analyses, and resistance selections across graded parasite inocula (minimum inoculum of resistance, MIR). Our findings provide a framework for distinguishing potential antiparasitic leads from compounds better suited to mechanistic investigation or medicinal-chemistry optimization.

## Results and Discussion

The ReFRAME library was plated in 384-well format and tested for growth inhibition of *T. gondii* isolate Tg68 *in vitro* using a single concentration of 10 µM. Growth inhibition was monitored for both tachyzoites and bradyzoites induced by alkaline stress, or glucose restriction with glutamine, following a recently published protocol ^9^. We choose the Tg68 line because exhibits slower *in vitro* replication of tachyzoites cultured in Human Foreskin Fibroblast cells (HFF), and undergoes permissive bradyzoite development under stress conditions with little breakthrough of tachyzoite growth.^9, 29^ From the primary screen, 160 compounds that inhibited parasite growth ≥ 80% in tachyzoites and bradyzoites were defined as “initial hits” and plotted as a VENN diagram (Figure 1A). To eliminate toxic compounds, 61 compounds with CC_50_ ≤ 5 µM were removed based on publicly available cytotoxicity data for HEK293T cells (Human Embryonic Kidney 293 with the SV40 large T-antigen) (https://reframedb.org/). The remaining 99 compounds were further evaluated using *in vitro* dilution series to monitor parasite growth inhibition (Tg68 EC_50_) and host cellular growth inhibition using Hepatocellular Grade 2 (HepG2) cells to determine CC_50_ values (Figure 1A). From these 99 compounds, 15 top hit compounds were identified based on Tg68 EC_50_ ≤ 1.0 µM and HepG2 CC_50_ vs Tg68 EC_50_ selectivity index (SI) ≥ 20-fold (Figure 1A). These 15 compounds were then sourced from commercial vendors or through collaboration (Table S1), and *in vitro* activity was reconfirmed for TgMe49 tachyzoites, as well as *in vitro* differentiated Tg68 bradyzoites generated by alkaline or glutamine-induction (Figure 1A). We also examined their activity on bradyzoites of the TgMe49EW strain that were isolated from cysts of the brains of chronically infected mice (referred to as *ex vivo* bradyzoites) using both liberated and intact cysts (Figure 1A).^9, 18, 19^ Finally, we evaluated their potential for developing resistance using a minimum inoculum of resistance (MIR) assay as described previously (Figure 1A).^30^ A previously described inhibitor of parasite phenylalanine tRNA synthetase BRD7929,^19^ atovaquone, which inhibits the bc1 complex^31^ and pyrimethamine that inhibits DHFR^32, 33^, were used as controls.

**Figure 1:**
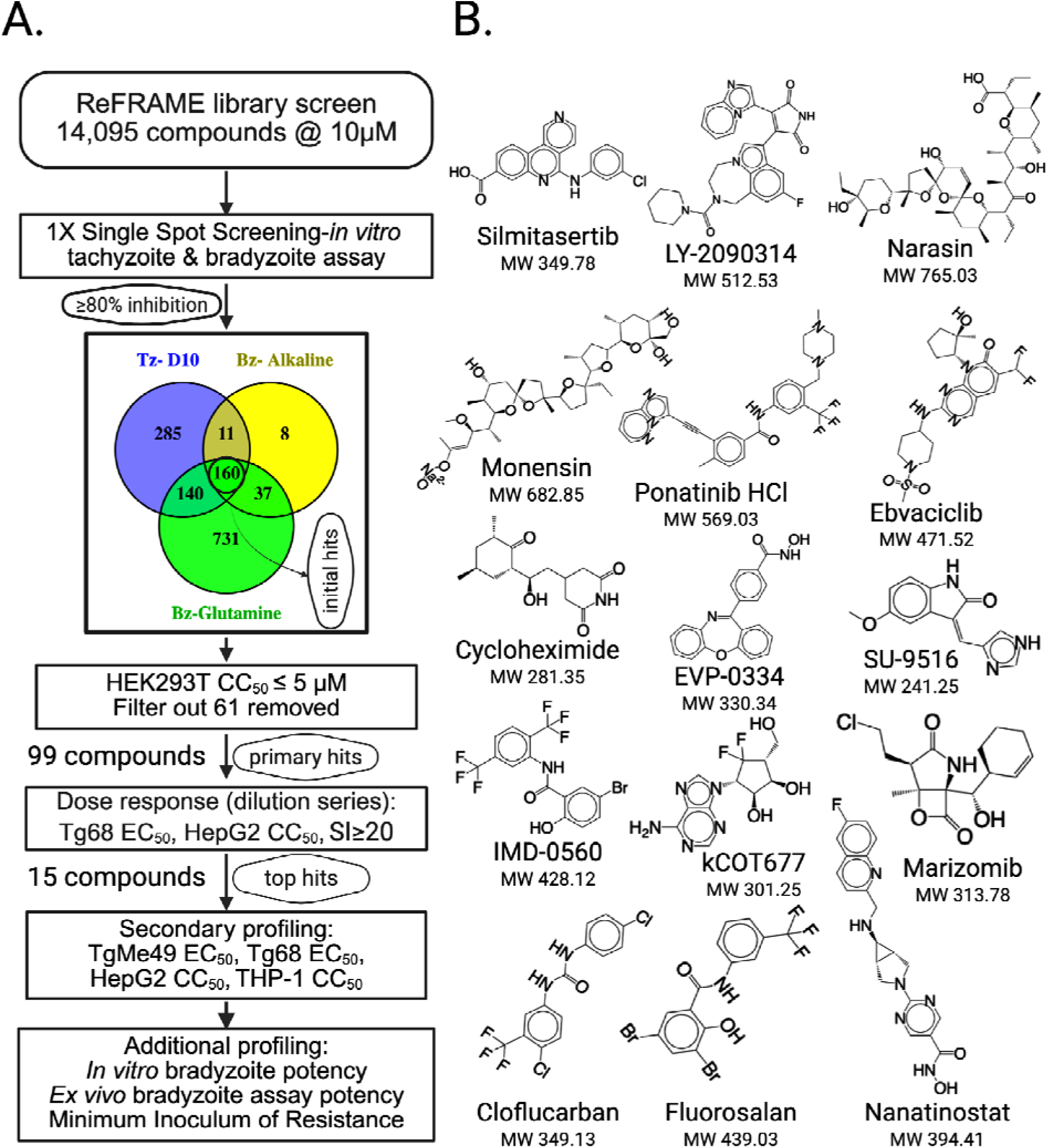
ReFRAME screening workflow, hit prioritization, and chemical structures of prioritized anti-*Toxoplasma gondii* compounds. A. The ReFRAME library consisting of 14,095 compounds was screened at 10 µM for growth inhibition of Tg68 tachyzoites and bradyzoites ^9^, yielding 160 hits with ≥ 80% growth inhibition. After exclusion of 61 cytotoxic compounds based on HEK293T CC_50_ ( ≤ 5 µM), 99 primary hits underwent dose–response testing (Tg68 EC_50_, HepG2 CC_50_). Fifteen highly potent compounds with a selectivity index (SI, host cell CC_50_/Tg68EC_50_) ≥ 20 were secondarily profiled for TgME49 EC_50_, and THP-1 CC_50_. Additional profiling included *in vitro* bradyzoite, *ex vivo* bradyzoite inhibition, and minimum inoculum of resistance assessment. B. Chemical structures and molecular weights of the top-15 hits.

The structures and molecular weights of the top hits are shown in Figure 1B and the results of secondary profiling of the tops hits are summarized in Table 1. Most compounds showed low-micromolar to sub-micromolar activity against tachyzoites from Tg68 and TgME49 strains. Notably, LY-2090314, narasin, monensin, marizomib, and nanatinostat exhibited excellent potency (EC_50_ ≤ 0.050 μM). In particular, LY- 2090314 showed a favorable *in vitro* selectivity profile, with a TgME49Fluc EC_50_ of 0.017 μM and Tg68 EC_50_ of 0.045 μM and selectivity indexes of 1,105 and 1,384 in HepG2 and differentiated THP-1 cells, respectively (Table 1). Narasin and monensin were the most potent compounds overall, with EC_50_ values of 0.001–0.004 μM in both tachyzoite and bradyzoite assays that was accompanied by low host-cell toxicity (Table 2). Monensin also displayed excellent selectivity in both host-cell lines (SI = 1,105 in HepG2 and 245 in THP-1 cells), whereas narasin showed moderate SIs of 925 and 493, respectively (Table 1). As expected, cycloheximide, despite nanomolar antiparasitic activity, was also highly cytotoxic (Table 1).^34^ Marizomib and nanatinostat retained potent antiparasitic activity but showed substantially lower selectivity in differentiated THP-1 cells (SI = 5 and 12, respectively), indicating a narrower selectivity window (Table 1). Together, these data identify LY-2090314, narasin, and monensin as the most potent and selective hits *in vitro* (Table 1). Several compounds showed condition- dependent activity in *in vitro* bradyzoite assays, including silmitasertib, ponatinib, EVP- 0334, and Marizomib which were 4−7-fold more active under glutamine-induction conditions than under alkaline-induction conditions, suggesting that parasite metabolic or differentiation-associated states may influence compound susceptibility.

**Table 1:** *In vitro* Potency, Stage-Specific Activity, and Selectivity Indexes of Top Hits^a^.

| **Compound** | **Tg68Fluc EC<sub>50</sub><br>(Tachyzoite)** | **TgME49Fluc EC<sub>50</sub><br>(Tachyzoite)** | **Tg68nLuc EC<sub>50</sub><br>(Alkaline)** | **Tg68nLuc EC50<br>(Glutamine)** | **HepG2<br>CC<sub>50</sub>** | **SI<sup>b</sup>** | **THP-1<sup>c</sup><br>CC<sub>50</sub>** | **SI** |
|---|---|---|---|---|---|---|---|---|
| Silmitasertib | 0.305 | 0.503 | 9.304 | 1.787 | 9.940 | 20 | 5.220 | 10 |
| LY-2090314 | 0.045 | 0.017 | 0.027 | 0.018 | 18.790 | 1105 | 23.530 | 1384 |
| Narasin | 0.001 | 0.004 | 0.003 | 0.002 | 3.700 | 925 | 1.970 | 493 |
| Monensin | 0.002 | 0.002 | 0.002 | 0.003 | 2.210 | 1105 | 0.490 | 245 |
| Ponatinib-HCl | 0.051 | 0.042 | 0.468 | 0.118 | 0.290 | 7 | 1.370 | 33 |
| Ebvaciclib | 0.125 | 0.088 | 1.577 | 0.598 | 3.730 | 42 | 4.530 | 51 |
| Cycloheximide | 0.025 | 0.005 | 0.080 | 0.152 | 2.500 | 500 | 1.860 | 372 |
| EVP-0334 | 0.056 | 0.044 | 0.629 | 0.157 | 4.050 | 92 | 0.920 | 21 |
| SU-9516 | 0.192 | 0.169 | 0.905 | 0.546 | 7.930 | 47 | 6.970 | 41 |
| IMD-0560 | 0.185 | 0.145 | 0.112 | 0.048 | 5.590 | 39 | 1.820 | 13 |
| kCOT677 | 0.087 | 0.057 | 0.211 | 0.177 | 6.540 | 115 | 4.830 | 85 |
| Cloflucarban | 0.350 | 0.450 | 0.549 | 0.541 | 10.500 | 23 | 6.140 | 14 |
| Fluorosalan | 1.815 | 1.611 | 1.790 | 0.747 | 11.290 | 7 | 10.380 | 6 |
| Marizomib | 0.007 | 0.004 | 0.027 | 0.004 | 0.560 | 140 | 0.020 | 5 |
| Nanatinostat | 0.010 | 0.005 | 0.091 | 0.148 | 0.680 | 136 | 0.060 | 12 |
| BRD7929 | 0.146 | 0.040 | 0.230 | 0.070 | 7.720 | 193 | 3.000 | 75 |
| Atovaquone | 0.286 | 0.043 | 0.881 | 0.057 | 12.180 | 283 | 15.310 | 356 |

<sup>a</sup>Values in *µ*M, average of three or more biological replicates. <sup>b</sup>SI = selectivity index (Host cell CC<sub>50</sub>/EC<sub>50</sub> TgME49Fluc). <sup>c</sup>THP-1 cells were differentiated as described in the Experimental Methods section. BRD7929 and atovaquone were included as controls.

**Table 2:** A summary of publicly available information on Drug Targets, Target Diseases, Trial Phases, and Safety Profiles of Top Hits.

| Compound | Target / Class | Human status | Takeaway and References |
|---|---|---|---|
| Silmitasertib (CX-4945) | CK2 inhibitor | Investigational; evaluated in Phase I/II oncology studies, including PBTC-053 in recurrent SHH medulloblastoma | Clinical-stage CK2 inhibitor; not FDA approved<sup>43, 44</sup> |
| LY-2090314 | GSK-3 inhibitor | Phase 1 completed (NCT01287520); limited clinical activity reported in AML | Clinical-stage exposure data available; no approved indication<sup>45-47</sup> |
| Ponatinib-HCl (ICLUSIG) | BCR-ABL / multi-TKI | FDA-approved (CML/Ph+ ALL) | Boxed warnings: arterial occlusion/VTE/HF/hepatotoxicity<sup>48-51</sup> |
| Ebvaciclib (PF-06873600) | CDK2/4/6 inhibitor | Phase 1/2a terminated (business decision; not safety) | Human data exist; program stopped<sup>52, 53</sup> |
| Narasin | Polyether ionophore coccidiostat | Not used in human medicine | Veterinary feed additive / non-translatable<sup>37, 54</sup> |
| Monensin | Polyether ionophore | Human toxicity reported (incl. fatal ingestion case) | Toxicity barrier; not human therapeutic<sup>55, 56</sup> |
| Cycloheximide | Eukaryotic protein synthesis inhibitor | Research tool; high toxicity risk | Inhibition of conserved host translation creates a major host-toxicity liability, not suitable for human therapy<sup>34, 57</sup> |
| EVP-0334 (FRM-0334) | HDAC inhibitor | Evaluated in a Phase IIa clinical study in FTD with GRN haploinsufficiency | Human-tested; PD/bioavailability issues reported<sup>58</sup> |
| SU-9516 | CDK inhibitor (CDK2; also listed CDK5) | Experimental; no clinical trials listed on DrugBank | Tool compound / preclinical oncology use<sup>40, 59</sup> |
| IMD-0560 | IKKβ/NF-κB-pathway inhibitor | Preclinical (xenograft models) | Anti-inflammatory/anti-tumor signaling inhibitor<sup>42, 60</sup> |
| kCOT677 | Viral RdRp + SAH hydrolase inhibition (nucleoside analog), Identity/target requires verification | Preclinical (no human data) | Dual-targeting antiviral, early-stage studies (No published data, information acquired through personal communication) |
| Cloflucarban | Halogenated carbanilide antiseptic/disinfectant agent | Topical antiseptic | Used in antimicrobial soaps/deodorants; not a systemic therapeutic candidate<sup>61</sup> |
| Fluorosalan | Halogenated salicylanilide antimicrobial | Biocidal/antimicrobial chemical | not established as therapeutic drug<sup>38</sup> |
| Marizomib | Irreversible pan-proteasome inhibitor | Evaluated in Phase I/II glioblastoma studies and later Phase III GBM study | CNS exposure is supported, but neurologic/CNS adverse events limit development<sup>62-65</sup> |
| Nanatinostat | Class-I selective HDAC inhibitor | Phase Ib/II evaluation with valganciclovir in relapsed/refractory EBV-positive lymphoma | Clinical responses reported; nausea and cytopenias are key adverse events; further Phase II evaluation is ongoing<sup>55, 66, 67</sup> |

Available public information for the 15 top hits, including annotated targets, prior clinical development, and reported safety considerations are summarized in Table 2. Several compounds have previously undergone clinical evaluation, and ponatinib-HCl is approved for treatment of BCR-ABL–positive malignancies.^35^ However, ponatinib-HCl carries FDA box warnings for arterial occlusive events, venous thromboembolic events, heart failure, and hepatotoxicity. Additionally, narasin and monensin are veterinary polyether ionophores with substantial toxicity^21, 36–38^ and cycloheximide is a eukaryotic translation inhibitor with substantial host toxicity.^34, 39^ These liabilities limit the potential for repurposing of these candidates for therapeutic treatment of toxoplasmosis, despite their potent activity on the parasite. In contrast, SU-9516, IMD-0560, and kCOT677 represent early-stage starting points for exploring mechanism of action.^40–42^

Fixed-ratio isobologram analyses were performed using TgME49Fluc tachyzoites to assess interactions between silmitasertib, ponatinib-HCl, EVP-0334, or marizomib and previously identified inhibitors atovaquone, BRD7929, or pyrimethamine (Figure 2, Figure S1). Most combinations produced mean ΣFIC_50_ (the Sum of Fractional Inhibitory Concentrations at 50% inhibition) values below 1.0 and were classified as additive.^68^ All four compounds showed reproducible additive interactions with atovaquone, with mean ΣFIC_50_values of 0.86 ± 0.02 for silmitasertib, 0.95 ± 0.01 for ponatinib, 0.84 ± 0.14 for EVP-0334, and 0.86 ± 0.08 for marizomib (Figure 2). Similarly, combinations with BRD7929 were additive overall, with marizomib–BRD7929 showing the lowest mean value (0.73 ± 0.06). Silmitasertib–pyrimethamine produced the lowest and most reproducible ΣFIC_50_ value among the pyrimethamine combinations (0.68 ± 0.01), followed by ponatinib–pyrimethamine (0.75 ± 0.10) and marizomib–pyrimethamine (0.75± 0.03). In contrast, EVP-0334–pyrimethamine yielded mean ΣFIC_50_ values above 1.0 in both experiments (1.24 and 1.13), resulting in an overall indifferent interaction (1.19 ± 0.08). The reference-drug combinations were also predominantly additive, with mean ΣFIC_50_ values of 0.84± 0.10 for atovaquone–BRD7929 and 0.84± 0.18 for atovaquone–pyrimethamine (Figure S1 A-B). The BRD7929–pyrimethamine combination was near the additivity boundary and was classified as indifferent overall (1.04± 0.12) (Figure S1 C). These findings indicate that the prioritized hits likely have mechanistically distinct anti-*Toxoplasma* reference compounds without evidence of antagonism.^69^

**Figure 2.**
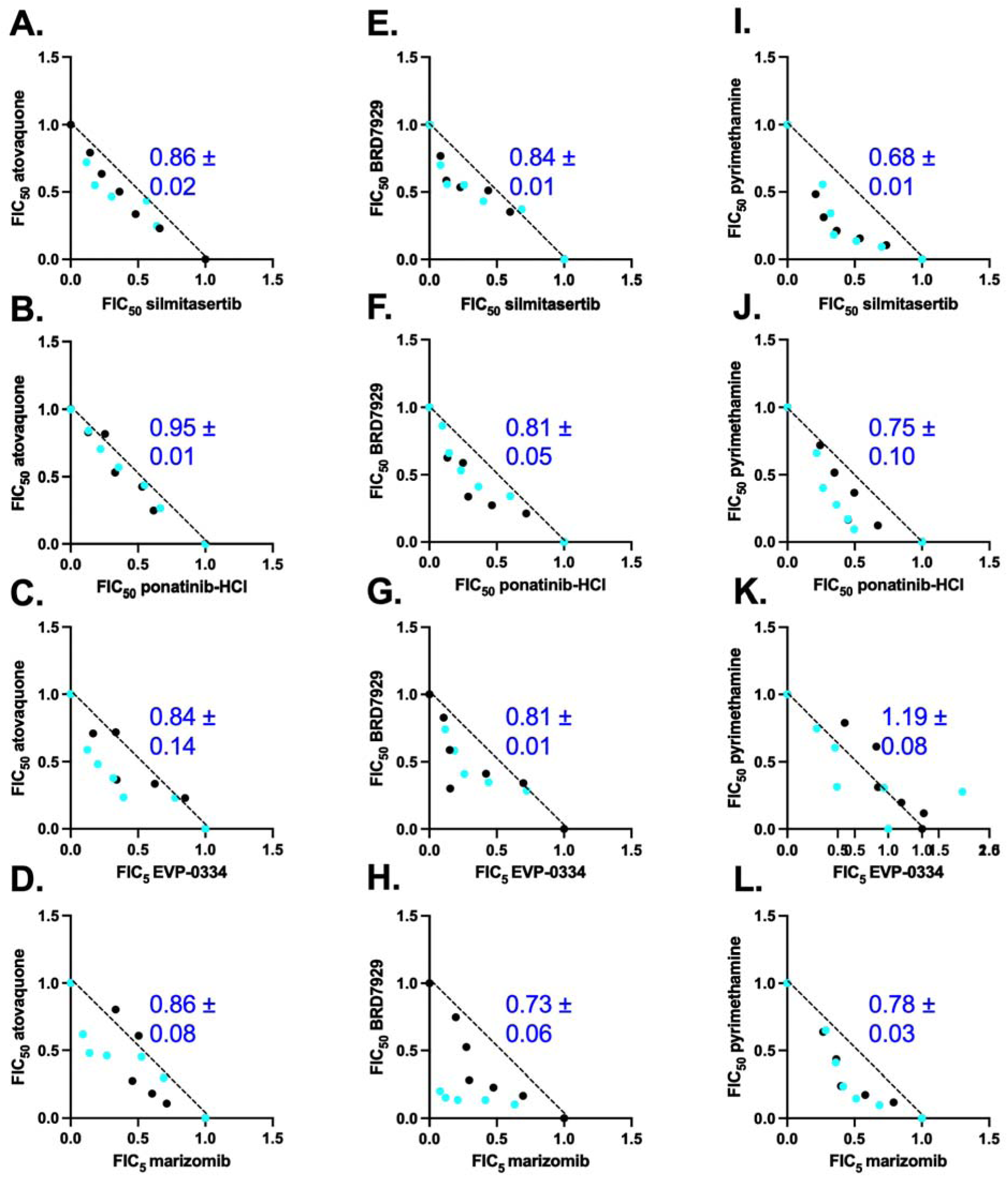
Top hits show predominantly additive interactions with reference anti-*Toxoplasma* compounds. Fixed-ratio drug-combination assays were performed with select compounds combined with atovaquone (A-D), BRD7929 (E-H), or pyrimethamine (I-L). Drug interactions were quantified by isobologram analysis and expressed as the Fractional Inhibitory Concentration index at 50% growth inhibition (FIC_50_) and theΣFIC_50_ values (blue text) represent the average ± S.D. from two independent biological replicates.

To determine whether the top hit compounds also exhibit activity against the chronic bradyzoite stage, they were evaluated for inhibition of Tg49EW bradyzoites isolated from brain cysts of chronically infected CBA/CaJ mice (Figure 3A).^9, 18, 19, 70^ We compared their activity against liberated bradyzoites treated for 4 h to bradyzoites within intact tissue cysts treated for 24 h (Figure 3A). After treatment, liberated bradyzoites were allowed to form plaques under either continuous compound exposure or no compound (washout) conditions. Under continuous 3X EC_90_ compound exposure, all tested compounds clearly suppressed plaque formation, although this may reflect inhibition of tachyzoite growth rather than killing of bradyzoites (Figure 3B). Although all of the compounds were able to inhibit plaque formation when used continuously, the washout arm showed substantial differences in the durability of inhibition (Figure 3B). Among the top hits, marizomib, ponatinib, and monensin showed the greatest activity within 4 h against liberated bradyzoites, with plaque formation remaining absent or nearly absent after washout. In contrast, nanatinostat, EVP-0334, cycloheximide, ebvaciclib, and silmitasertib showed little-to-no-inhibition after compound removal, with residual plaque formation generally approaching control levels (Figure 3B). The compounds IMD-0560, SU-9516, cloflucarban, fluorosalan, kCOT677, narasin, and LY- 2090314 displayed intermediate post-washout effects, with variable recovery across biological replicates (Figure 3B). The intact-cyst assay faced the additional barrier of compound penetration into, or activity within, the bradyzoite-containing tissue cyst (Figure 3C). Although all compounds were again active when exposed continuously, they fared substantially less well when only used on intact cysts for 24 h. Marizomib, monensin, narasin, and LY-2090314 showed the strongest sustained effects after washout, with no or minimal residual plaque formation (Figure 3C). Nanatinostat and SU-9516 also retained substantial activity, although recovery was observed in some replicates (Figure 3C). The remaining compounds ponatinib, ebvaciclib, silmitasertib, EVP-0334, IMD-0560, kCOT677, cloflucarban, fluorosalan, and cycloheximide showed limited activity against intact cysts (Figure 3C). Of the control compounds, BRD7929 proved most effective in short term 4 h treatment and showed partial activity in the 24 h treatment of intact cysts, as did atovaquone (Figure 3B). Collectively, these findings identify marizomib, monensin, narasin, and LY-2090314 as the most compelling compounds based on their ability to prevent outgrowth after 24 h exposure to intact cysts.

**Figure 3:**
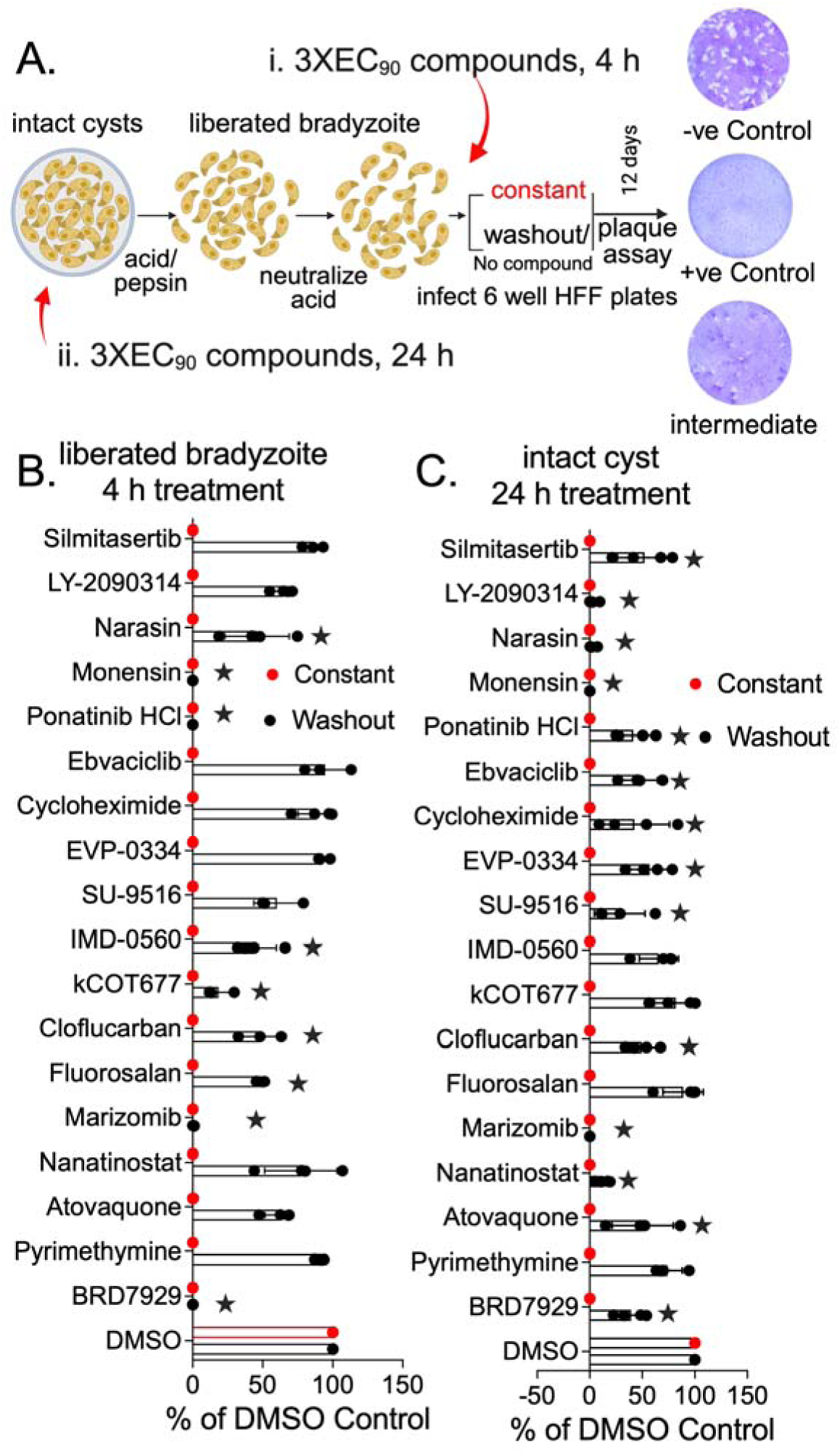
*Ex vivo* activity of top hits against liberated bradyzoites and intact tissue cysts under continuous-treatment and washout conditions. (A) Assay workflow. Brain cysts isolated from chronically infected mice were: 1) disrupted by pepsin to liberate bradyzoites and treated for 4 h treatment, or 2) treated intact for 24 h before pepsin disruption. Compounds were tested at 3X EC_90_, followed by 12 days of plaque development under continuous-treatment or washout conditions. (B, C) Inhibition of plaque formation following treatment of liberated bradyzoites (B) or intact cysts (C), relative to DMSO controls. Data represent mean ± SD from ≥ 3 independent biological replicates. Statistical comparisons with DMSO controls used one-way ANOVA followed by Dunnett’s multiple comparisons (*P* < 0.05); variance homogeneity was assessed using Brown–Forsythe and Bartlett’s tests.

To assess the propensity for resistance emergence, a minimum inoculum for resistance (MIR) assay was performed using TgME49-Fluc Clone B6 tachyzoites parasites at three different inocula (10^6^, 10^7^, and 10^7^) under continuous compound pressure at 3X EC_90_ for 30 days.^30^ This assay evaluates whether resistant parasites emerge from progressively larger parasite populations, thereby providing an experimental measure of their barrier to resistance emergence.^30^ Compounds SU- 9516, monensin, narasin, IMD-0560, EVP-0334, and marizomib completely suppressed detectable parasite growth in all three replicate flasks (Figure 4A). In contrast, LY- 2090314 and nanatinostat showed inoculum-dependent emergence of resistant parasites and ebvaciclib resulted in parasite outgrowth at all inocula tested (Figure 4A). Despite the reproducible emergence of parasites during drug exposure, the isolated ebvaciclib-resistant populations displayed only modest shifts in susceptibility, with EC_50_ values increased by approximately 1–7-fold compared with WT parasites (Figure 4C). Nanatinostat displayed an intermediate phenotype (Figure 4D). No parasite growth was detected in flasks initiated with 10^6^ or 10^7^ parasites, whereas parasite outgrowth occurred in all three flasks initiated with 10^8^ parasites. Nanatinostat-resistant parasites showed 4–6-fold increase in EC_50_ relative to the parental (WT) parasite line (Figure 4D). Two flasks inoculated with 10^8^ parasites resulted in resistance to LY-2090314 and these populations showed a 26–29-fold increase in EC_50_ relative to the parental TgME49-Fluc (WT) parasite line (Figure 4B). The emergence of mutants that were resistant to LY- 2090314 is consistent with recent work identifying parasite glycogen synthase kinase 3 (TgGSK3) as a target of LY-2090314.^71^ Point mutations in TgGSK3 have been shown to reduce LY-2090314 susceptibility in *T. gondii*, supporting the possibility that the high- level resistance detected here may result from target-based genetic changes. Collectively, our findings indicate that LY-2090314 and Nanatinostat have measurable, but relatively high, resistance barriers under the conditions tested.

**Figure 4.**
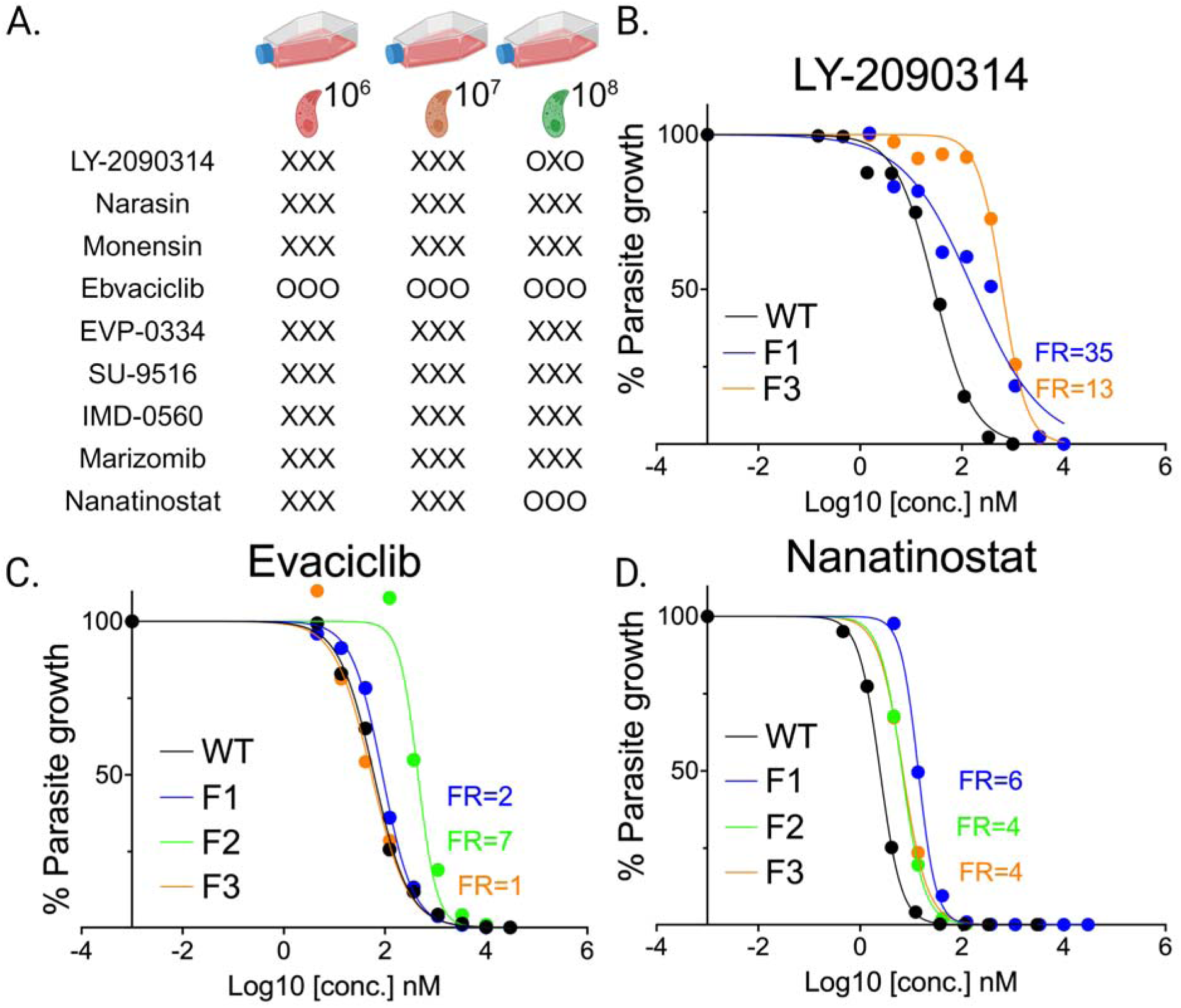
Profiling resistance via Minimum Inoculum for Resistance (MIR) assay. (A) Schematic summary of outcomes for compounds tested with different parasite inocula (10^6^ to 10^8^) in this study: X = No parasite growth in 30 days. O = parasite growth in 30 days. Each X or O represents one flask. (B) EC_50_ inhibition curves for wild type (WT) vs. two LY-2090314-resistant parasite populations: F1, and F3. (C) EC_50_ inhibition curves WT vs three ebvaciclib-resistant parasite populations: F1, F2, and F3. (D) EC_50_ inhibition curves WT vs three nanatinostat-resistant parasite populations: F1, F2, and F3. All EC_50_ values are presented as the mean of two or more biological replicates with two technical replicates per assay. Source data are provided in Table S3. FR = fold resistance; WT = parental TgME49-Fluc parasite.

To identify genetic changes associated with LY-2090314 resistance, we performed targeted sequencing of *TgGSK3* (ToxoDB *TGME49_265330*), which has previously been shown to be the target of LY-2090314.^71^ Resistant populations were cloned by limiting dilution, and four clones (C1–C4) from each of two independent populations were analyzed by full-length *TgGSK3* amplicon sequencing (Figure 5A). All eight clones carried the same nonsynonymous substitution, L53V, which was also detected in both resistant bulk populations (F1 and F3). LY-2090314 is an ATP-competitive inhibitor of mammalian GSK-3, and recent genetic, biochemical, and structural studies identify TgGSK3 as its principal parasite target.^71, 72^ In the published study, TgGSK3 substitutions G54D, H129D, and T133M, reconstructed in the susceptible parental background, reduced LY-2090314 susceptibility by approximately 2.9-, 4.5-, and 4.8- fold, respectively.^71^ In contrast, the resistant populations generated here exhibited a 26– 29-fold increase in LY-2090314 EC_50_ relative to the parental line (Figure 4B). Thus, L53V expands the set of resistance-associated substitutions and further supports TgGSK3a as the target of LY-2090314 action.

**Figure 5.**
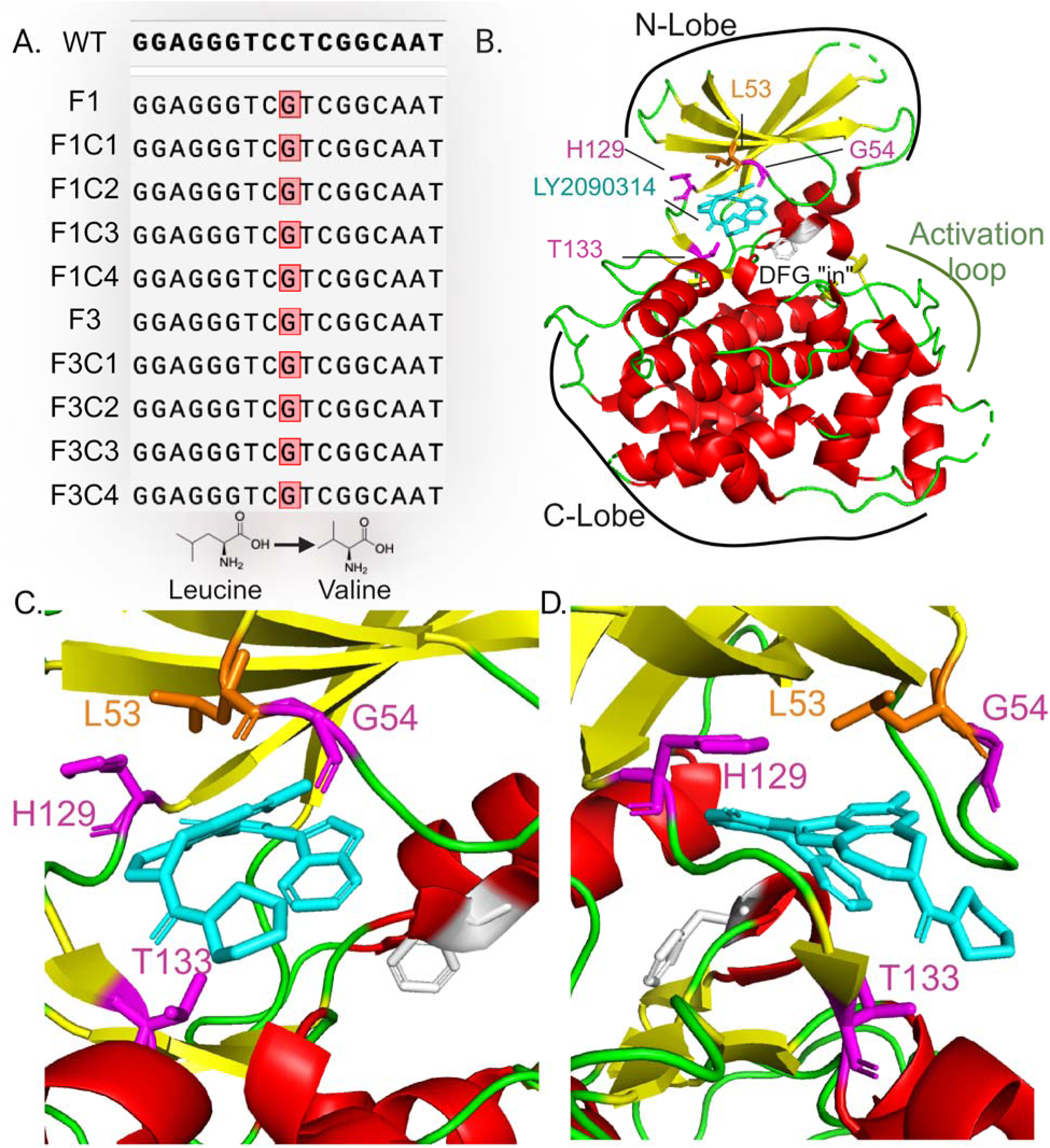
Identification and structural mapping of the TgGSK3 L53V substitution in LY-2090314- resistant *Toxoplasma gondii*. (A) Resistance mutations identified in *TgGSK3* from four limiting-dilution clones (C1–C4) from each of two independently selected resistant populations (F1 and F3) identified a CTC-to-GTC codon change, resulting in an L53V substitution in all eight clones. (B) The cartoon representation shows alpha helices in red, beta-strands in yellow and loops in green. The LY2090314 is colored cyan and displayed in a stick representation. F196 (from the DFG motif) is displayed in grey. Previously reported G54, H129, T133 mutant sites are displayed in magenta, while the new mutant from current study L53 is displayed in orange. N-and C-lobes are indicated. C. Enlargement of the LY2090314 binding site. Using the same color scheme as in B. Side chains involved in the compound interactions are displayed, with resistance conferring residues shown. D. Rotation of the C representation by 90°. Structural representations were generated in PyMOL (version 3.1.4.1) using the published crystal structure deposited in the Protein Data Bank (PDB ID: 9HVX).

Structural mapping placed L53 immediately adjacent to G54 in the glycine-rich loop of the TgGSK3 N-lobe (Figure 5B). The corresponding position in human GSK3 is isoleucine rather than leucine, indicating a local sequence difference between the parasite and human kinases.^71^ Although both residues are hydrophobic, their different side-chain branching provides a potential basis for differences in host vs. parasite selectivity. In the published TgGSK3–LY-2090314 co-crystal structure (PDB ID 9HVX), the compound binds the ATP pocket as a type I inhibitor; its maleimide core forms hinge-backbone hydrogen bonds with E128 and I130, whereas T133 makes a direct side-chain interaction with the ligand.^71^ G54D and H129D are proposed to reduce inhibitor binding indirectly by altering local backbone geometry, whereas T133M disrupts a direct interaction and introduces steric interference.^71^ By analogy, the conservative L53V substitution may alter inhibitor susceptibility through changes in local hydrophobic packing or conformational dynamics of the L53–G54 segment (Figure 5C-D).

### Conclusion

Screening of the ReFRAME library identified 15 potent anti-*Toxoplasma gondii* hits that were active against tachyzoites and bradyzpoites. LY-2090314, narasin, and monensin showed the greatest *in vitro* potency and selectivity, whereas marizomib, monensin, narasin, and LY-2090314 demonstrated durable activity against intact-cyst treated for only 24 hours. The top hist were largely additive with atovaquone, BRD7929, or pyrimethamine suggesting they have unrelated mechanisms of action. MIR assays distinguished compounds with different apparent barriers to resistance. LY-2090314- resistant parasites emerged only from the highest inoculum and contained a consistent *TgGSK3* L53V substitution, supporting previous findings that TgGSK3 is the parasite target. Overall, these findings identify chronic-stage-active compounds that require further evaluation of parasite target, *in vivo* PK and efficacy, and host-toxicity before further development.

## Experimental Methods

### Compounds and Liquid Handling

Initially, the ReFRAME library (14,095 compounds), along with positive and negative controls, was dispensed by our collaborator at Calibr, Scripps Research (La Jolla, CA), in single-spot format into 384-well plates for primary screening of percent growth inhibition against tachyzoite and in vitro–induced bradyzoite stages. Compound plates stored at −80 °C until use. Primary screening assays (one tachyzoite and two bradyzoite formats- in alkaline and glutamine induction media) were performed in a single biological replicate. Host and parasite cell seeding, media exchanges, compound transfers, and luciferase reagent additions were carried out at the semi-automated High-Throughput Screening Center at Washington University School of Medicine to ensure consistency and reproducibility across replicates. Calibr also dispensed the 99 primary hits in 3-fold dilution series to determine Tg68 EC_50_ values and HepG2 CC_50_ in duplicate. For top-hit validation, an assay control compound (BRD7929) was provided by our collaborator at Calibr, and two additional control compounds—atovaquone (Sigma, A7986) and pyrimethamine (Sigma, 46706)—were purchased from commercial sources. All follow-up assays were performed manually in-house using 96-well plates.

### *In Vitro* Assays for Host cell cytotoxicity

A HEK293 cytotoxicity dataset generated by collaborators at Calibr, Scripps Research (La Jolla, CA) was obtained from the publicly available ReFRAME database (https://reframedb.org/). Compounds were pre- spotted (up to 50 nL) into white, solid-bottom, tissue culture-treated 1536-well plates. HEK293 cells were dispensed at 1,500 cells/well in 4 μL of DMEM supplemented with 5% FBS and incubated at 37°C with 5% CO_₂_for 72 h. After incubation, 4 μL of CellTiter- Glo reagent (Promega#G7571) was added per well, and luminescence was measured. This dataset was used to eliminate 61 compounds with CC_₅₀_ values ≤5 μM from the initial set of 160 hits (Figure1). Remaining 99 primary hits were further evaluated for cytotoxicity against HepG2 cells (ATCC HB-8065). All host cell lines were confirmed to be mycoplasma-free using the e-Myco Plus kit (Intron Biotechnology). Compounds were diluted to 2X working concentrations, and a 10-point, 3-fold serial dilution series was prepared. HepG2 cells were seeded at a density of 1X10^3^ cells/well in 40 μL of medium to produce sub-confluent monolayers for a 72 h assay. Compounds were added 6 h post-seeding (80 μL final volume, 0.05% DMSO final concentration), and plates were incubated at 37°C in 5% CO_2_. Seventy-two hours after compound addition, culture volumes were reduced to 20 μL, followed by the addition of an equal volume of CellTiter-Glo Luminescent Cell Viability Assay reagent (Promega, G7571). Luminescence was measured using a BioTek Cytation 3 equipped with Gen5 software (v3.08). Each assay included two technical replicates and was repeated in at least two independent biological experiments. During secondary screening, cytotoxicity profiling was expanded to include the human monocytic cell line THP-1 (ATCC TIB-202), maintained according to ATCC recommendations. Prior to compound treatment, THP-1 cells were differentiated into macrophage-like cells by incubation with 10 ng/mL phorbol 12-myristate 13-acetate (PMA) for 24 h. Statistical analyses were performed using GraphPad Prism 10. CC_50_ values were determined by nonlinear regression using the “log(inhibitor) versus normalized response – variable slope” model.

### *In vitro* Assays for tachyzoite and bradyzoite growth inhibition

Tg68 pTub1:Fluc, Tg68 pBAG1:nLuc and TgME49 pTub1:Fluc parasites used in this study were described previously.^9, 73^ Human Foreskin Fibroblasts (HFF-1 SCRC-1041, ATCC) cells were cultured in T175 flasks in D10 medium. Prior to plating, cells were detached by treatment with trypsin solution (0.5 g/L porcine trypsin and 0.2 g/L EDTA in Hank’s Balanced Salt Solution containing phenol red) for 5 min to generate a single-cell suspension. EC_50_ assays (half maximal effective concentration) were conducted in a 384-well plate format. Confluent HFF cells were seeded 4 days in advance using a Multidrop Combi dispensing 80 uL/well, while stirring the cell suspension at 350 rpm. The plates were stacked in a Cytomat rack on a level surface at room temperature until all plates were completed, and then they were placed in a 37°C incubator with 5% CO_2_. To minimize edge effects, only the inner 240 wells of each plate were utilized. Tachyzoite and bradyzoite growth inhibition assays were performed as previously described.^9^

### Drug Combination Isobologram assay

Drug interactions were evaluated using fixed- ratio isobologram analyses in two biological replicates. The following fixed ratios were prepared as starting concentrations: 1:0, 4:1, 2:1, 1:1, 1:2, 1:4, and 0:1, to assess interactions between silmitasertib, ponatinib-HCl, EVP-0334, or marizomib and atovaquone, BRD7929, or pyrimethamine. The concentration of each drug alone (1:0 and 0:1 ratios) was set at approximately 16X EC_50_ value. TgME49Fluc tachyzoites were seeded at 10^4^ parasites per well and exposed to twofold serial dilutions of each drug combination ratio. Fractional inhibitory concentration at 50% inhibition (FIC_50_) values were calculated using the equation: FIC_50_= IC_50_ of drug in combination / IC_50_ of drug alone. For each drug pair, the FIC_50_ values of the two compounds were plotted on an XY graph to generate isobolograms. The sum of the fractional inhibitory concentrations (ΣFIC_50_) was calculated for each ratio, and the mean ΣFIC_50_was determined by averaging the values obtained from the 4:1, 2:1, 1:1, 1:2, and 1:4 combination ratios. Most combinations produced m Mean ΣFIC_50_ (the Sum of Fractional Inhibitory Concentrations at 50% inhibition) for combination is classified as additive when ΣFIC_50_ > 0.5 ≤ 1.0; synergy ≤ 0.5 and antagonism >2.0.^68^

### *Ex-vivo* Bradyzoite differentiation assay

CBA/CaJ mice (Strain 517#000654) from Jackson Laboratory were housed at the Washington University School of Medicine in an approved facility. All animal studies adhered to ethical guidelines approved by the National Institute of Animal Care and Use Committee. Production of chronically infected mice and isolation of tissue cysts were conducted using previously described protocols.^9, 19^ Briefly, CBA/CaJ mice were infected by oral gavage with 5–10 TgME49EW tissue cysts.^74^ At 1–2 months post- infection, brains from TgME49-EW-infected mice were harvested, homogenized, and tissue cysts were purified using Percoll gradients.^9, 18, 19, 70^ Bradyzoites were liberated by acid-pepsin digestion and neutralized with 94 mM Na_2_CO_3_ buffer. Liberated bradyzoites were distributed into duplicate 6-well plates containing D10 medium. Each plate included DMSO (0.1%) as a negative control, BRD7929 (0.5 μM), and pyrimethamine (2.0 μM) as positive controls. Test compounds were evaluated at EC_90_ and 3XEC_90_ concentrations based on their tachyzoite growth inhibition values (Table 2). After 4 h of treatment, compounds were removed from one set of plates by PBS washes (3X) and replacement with compound-free medium. Plates were then incubated at 37°C with 5% CO_2_ for 12–14 days to allow plaque formation. Plaques were visualized by ethanol fixation and crystal violet staining and quantified using a Nikon Eclipse TS2 microscope with a 4X objective. Plaque counts from at least three biological replicates were normalized to the DMSO control and expressed as % survival. For intact cyst-treatment, ∼50 purified cysts were treated with compounds at 3XEC_90_ for 4 or 24 h at 37°C. Following treatment, cysts were digested with acid-pepsin to release bradyzoites and plated in the presence or absence of compound. After 12–14 days of incubation, plaques were quantified as described above.

### Assessment of resistance emergence by MIR assay

MIR assays were performed as previously described using nine top-hit compounds.^30^ Confluent HFF monolayers in T25, T75, and T175 flasks were infected with 10^6^, 10^7^, and 10^8^ TgME49-Fluc tachyzoites, respectively, and cultured in D10 medium containing 3XEC_90_ compound concentrations. Cultures were maintained for up to 30 days with fresh compound- containing medium replacement every 3–7 days. Parasites exhibiting growth were expanded and evaluated for changes in EC_50_ values using the luciferase growth inhibition assay described above. Resistant populations were cloned by limiting dilution.

### Mutation mapping of LY-2090314-resistant parasites

Genomic DNA was isolated and sequenced from eight independent LY-2090314 resistant clonal parasites derived from two resistant population (four clones per population) as previously described.^30^ Parasites were purified from infected HFF cultures by syringe lysis, filtration through a 3-µm membrane, and centrifugation. Genomic DNA was extracted using the DNeasy Blood & Tissue Kit (QIAGEN). The full-length *TgGSK3* gene (*ToxoDB* TgME49_265330) was amplified using PrimeSTAR GXL Premix (Takara Bio) with primers TgMe49_265330_FwPrimer CTCGCGGCAAGTTTCTTCAC, TgMe49_265330_RvPrimer GTGCTCGATGCGTGTCTCTA. Purified PCR products were subjected to long-read amplicon sequencing (Plasmidsaurus; ONT Rapid Barcoding Kit V14, PromethION platform). Sequence reads were aligned to the parental *TgGSK3* sequence and analyzed using SnapGene v8.2.3 to identify resistance-associated mutations.

### Structure models of the resistance mutations

Structural representations were generated in PyMOL version 3.1.4.1 using the published wildtype TgGSK3–LY-2090314 crystal structure (Protein Data Bank entry 9HVX). The kinase was displayed as a cartoon, with α-helices colored red, β-strands yellow, and loop regions green. LY- 2090314 was displayed as cyan sticks, and the DFG-motif residue F196 was shown as gray sticks. The previously reported LY-2090314 resistance-associated residues G54, H129, and T133 were displayed as magenta sticks, whereas L53, identified in the present study, was shown as orange sticks. The N- and C-lobes of the kinase were labeled, and close-up views of the ATP-binding pocket were generated in two orientations, including a 90° rotation. Structural views depict the wild-type TgGSK3–LY- 2090314 complex; no in silico mutagenesis, docking, molecular-dynamics simulation, or biochemical analysis of L53V was performed. Therefore, the structural consequences of L53V were not experimentally determined.

### AI-assisted organization of compound information

Perplexity.ai (https://www.perplexity.ai/; accessed June-September 2025; free online version) was used to organize preliminary content from accumulated dossier for Table 2. The author independently reviewed and revised the content and verified the information and references.

### Statistical analysis

Statistical comparisons were performed in Prism (GraphPad). Data were first analyzed for normal distribution and according to the outcome, were then non-parametric tests applied. Statistical test and P values are given in the figure legends.

## ASSOCIATED CONTENT

**\*sı Supporting Information**

Table S1: Sources and catalog numbers of the top 15 hit compounds.

Table S2: EC_₅₀_ values and fold resistance of WT and resistant populations.

Figure S1: Isobologram analysis of interactions among reference anti-*Toxoplasma* compounds in TgME49Fluc tachyzoites.

## Author Contributions

Conceptualization: T.U., L.D.S.; Data curation: T.U.; Formal analyses: T.U.; Investigation: T.U.; Supervision: L.D.S.; Writing original draft: T.U.; Writing review and editing: all authors.

## Notes

The authors declare no competing financial interest.

## Acknowledgments

We thank Jennifer Barks for cell culture support and assistance with research administration. We acknowledge Maxene Ilagan and Mike Prinsen of the Washington University High Throughput Screening Center for assay setup and data management, and Melissa Rosenthal, Washington Univ. Sch. of Medicine for assistance with isobologram assay design and analysis. We also thank Kaycie Morwood and colleagues at Calibr, a Division of Scripps Research, for compound management and plating support. The TgME49-EW strain was generously provided by Prof. Michael White (University of South Florida). This work was supported in part by NIH grant AI143857 and a Center for Drug Discovery pilot grant at Washington University (PJ00027513).

## Abbreviations used

DMSO: Dimethylsulfoxide
DHFR: Dihydrofolate Reductase
EDTA: Ethylenediaminetetraacetic acid
HFF: Human Foreskin Fibroblasts
HepG2: human hepatocellular carcinoma cell
THP-1: human monocytic tumor cell
HTS: High Throughput Screening
nLuc: Nano Luciferase
Fluc: Firefly Luciferase
PheRS: Phenylalanine tRNA synthetase
WT-: Wild type

## References

(1) Dubey, J. P. Advances in the life cycle of Toxoplasma gondii. Int J Parasitol 1998, 28 (7), 1019–1024.

(2) Zhao, X. Y.; Ewald, S. E. The molecular biology and immune control of chronic Toxoplasma gondii infection. J Clin Invest 2020, 130 (7), 3370–3380.

(3) Sullivan, W. J., Jr.; Jeffers, V. Mechanisms of Toxoplasma gondii persistence and latency. FEMS Microbiol Rev 2012, 36 (3), 717–733.

(4) Daher, D.; Shaghlil, A.; Sobh, E.; Hamie, M.; Hassan, M. E.; Moumneh, M. B.; Itani, S.; El Hajj, R.; Tawk, L.; El Sabban, M.;, et al. Comprehensive Overview of Toxoplasma gondii-Induced and Associated Diseases. Pathogens 2021, 10 (11), 1351.

(5) Wang, Z. D.; Liu, H. H.; Ma, Z. X.; Ma, H. Y.; Li, Z. Y.; Yang, Z. B.; Zhu, X. Q.; Xu, B.; Wei, F.; Liu, Q. Toxoplasma gondii Infection in Immunocompromised Patients: A Systematic Review and Meta-Analysis. Front Microbiol 2017, 8, 389.

(6) Konstantinovic, N.; Guegan, H.; Stajner, T.; Belaz, S.; Robert-Gangneux, F. Treatment of toxoplasmosis: Current options and future perspectives. Food Waterborne Parasitol 2019, 15, e00036.

(7) Dunay, I. R.; Gajurel, K.; Dhakal, R.; Liesenfeld, O.; Montoya, J. G. Treatment of Toxoplasmosis: Historical Perspective, Animal Models, and Current Clinical Practice. Clin Microbiol Rev 2018, 31 (4), e00057–00017.

(8) Alday, P. H.; Doggett, J. S. Drugs in development for toxoplasmosis: advances, challenges, and current status. Drug Des Devel Ther 2017, 11, 273–293.

(9) Uddin, T.; Xia, J.; Fu, Y.; McNamara, C. W.; Chatterjee, A. K.; Sibley, L. D. High- Throughput Repurposing Screen Reveals Compounds with Activity against Toxoplasma gondii Bradyzoites. ACS Infect Dis 2025, 11 (3), 600–609.

(10) Cerutti, A.; Blanchard, N.; Besteiro, S. The Bradyzoite: A Key Developmental Stage for the Persistence and Pathogenesis of Toxoplasmosis. Pathogens 2020, 9 (3),

(11) Maus, D.; Putrianti, E.; Hoffmann, T.; Laue, M.; Seeber, F.; Blume, M. Screening the MMV Pathogen Box reveals the mitochondrial bc(1)-complex as a drug target in mature Toxoplasma gondii bradyzoites. Elife 2026, 13,

(12) Smith, D.; Lunghi, M.; Olafsson, E. B.; Hatton, O.; Di Cristina, M.; Carruthers, V. B. A High-Throughput Amenable Dual Luciferase System for Measuring Toxoplasma gondii Bradyzoite Viability after Drug Treatment. Anal Chem 2023, 95 (2), 668–676.

(13) Sinai, A. P.; Watts, E. A.; Dhara, A.; Murphy, R. D.; Gentry, M. S.; Patwardhan, A. Reexamining Chronic Toxoplasma gondii Infection: Surprising Activity for a “Dormant” Parasite. Curr Clin Microbiol Rep 2016, 3 (4), 175–185.

(14) Kim, K. A Bradyzoite is a Bradyzoite is a Bradyzoite? Trends Parasitol 2015, 31 (12), 610–612.

(15) Ortiz, J. O.; Potter, A. K.; Benmerzouga, I. Novel Drug Targets for the Bradyzoite Form of Toxoplasma gondii. Res Rep Trop Med 2025, 16, 25–30.

(16) Murata, Y.; Sugi, T.; Weiss, L. M.; Kato, K. Identification of compounds that suppress Toxoplasma gondii tachyzoites and bradyzoites. PLoS One 2017, 12 (6), e0178203.

(17) Dittmar, A. J.; Drozda, A. A.; Blader, I. J. Drug Repurposing Screening Identifies Novel Compounds That Effectively Inhibit Toxoplasma gondii Growth. mSphere 2016, 1 (2), e00042–00015.

(18) Uddin, T.; Xie, H.; Mittal, P.; Ence, C. C.; Patil, S.; Sharma, A.; Melillo, B.; Chatterjee, A. K.; Sibley, L. D. Improved Bicyclic Pyrrolidine Analogues Inhibit Toxoplasma gondii Growth In Vitro and Cure Infection In Vivo. J Med Chem 2025, 68 (16), 17350–17363.

(19) Radke, J. B.; Melillo, B.; Mittal, P.; Sharma, M.; Sharma, A.; Fu, Y.; Uddin, T.; Gonse, A.; Comer, E.; Schreiber, S. L.;, et al. Bicyclic azetidines target acute and chronic stages of Toxoplasma gondii by inhibiting parasite phenylalanyl t-RNA synthetase. Nat Commun 2022, 13 (1), 459.

(20) Araujo, F. G.; Huskinson-Mark, J.; Gutteridge, W. E.; Remington, J. S. In vitro and in vivo activities of the hydroxynaphthoquinone 566C80 against the cyst form of Toxoplasma gondii. Antimicrob Agents Chemother 1992, 36 (2), 326–330.

(21) Couzinet, S.; Dubremetz, J. F.; Buzoni-Gatel, D.; Jeminet, G.; Prensier, G. In vitro activity of the polyether ionophorous antibiotic monensin against the cyst form of Toxoplasma gondii. Parasitology 2000, 121 (Pt 4) (4), 359–365.

(22) Maubon, D.; Bougdour, A.; Wong, Y. S.; Brenier-Pinchart, M. P.; Curt, A.; Hakimi, M. A.; Pelloux, H. Activity of the histone deacetylase inhibitor FR235222 on Toxoplasma gondii: inhibition of stage conversion of the parasite cyst form and study of new derivative compounds. Antimicrob Agents Chemother 2010, 54 (11), 4843–4850.

(23) Metaferia, B.; Cellmer, T.; Dunkelberger, E. B.; Li, Q.; Henry, E. R.; Hofrichter, J.; Staton, D.; Hsieh, M. M.; Conrey, A. K.; Tisdale, J. F.;, et al. Phenotypic screening of the ReFRAME drug repurposing library to discover new drugs for treating sickle cell disease. Proc Natl Acad Sci U S A 2022, 119 (40), e2210779119.

(24) Bernatchez, J. A.; Chen, E.; Hull, M. V.; McNamara, C. W.; McKerrow, J. H.; Siqueira-Neto, J. L. High-Throughput Screening of the ReFRAME Library Identifies Potential Drug Repurposing Candidates for Trypanosoma cruzi. Microorganisms 2020, 8 (4), 472.

(25) Janes, J.; Young, M. E.; Chen, E.; Rogers, N. H.; Burgstaller-Muehlbacher, S.; Hughes, L. D.; Love, M. S.; Hull, M. V.; Kuhen, K. L.; Woods, A. K.;, et al. The ReFRAME library as a comprehensive drug repurposing library and its application to the treatment of cryptosporidiosis. Proc Natl Acad Sci U S A 2018, 115 (42), 10750–10755.

(26) Riva, L.; Yuan, S.; Yin, X.; Martin-Sancho, L.; Matsunaga, N.; Pache, L.; Burgstaller-Muehlbacher, S.; De Jesus, P. D.; Teriete, P.; Hull, M. V.;, et al. Discovery of SARS-CoV-2 antiviral drugs through large-scale compound repurposing. Nature 2020, 586 (7827), 113–119.

(27) Kim, Y. J.; Cubitt, B.; Chen, E.; Hull, M. V.; Chatterjee, A. K.; Cai, Y.; Kuhn, J. H.; de la Torre, J. C. The ReFRAME library as a comprehensive drug repurposing library to identify mammarenavirus inhibitors. Antiviral Res 2019, 169, 104558.

(28) Maher, S. P.; Bakowski, M. A.; Vantaux, A.; Flannery, E. L.; Andolina, C.; Gupta, M.; Antonova-Koch, Y.; Argomaniz, M.; Cabrera-Mora, M.; Campo, B.;, et al. A drug repurposing approach reveals targetable epigenetic pathways in Plasmodium vivax hypnozoites. Elife 2025, 13,

(29) Xia, J.; Fu, Y.; Huang, W.; Uddin, T.; Sibley, L. D. Constitutive upregulation of transcription factors underlies permissive bradyzoite differentiation in a natural isolate of Toxoplasma gondii. mBio 2024, 15 (9), e0064124.

(30) Uddin, T.; Mittal, P.; Xie, H.; Melillo, B.; Sharma, A.; Chatterjee, A. K.; Sibley, L. D. Minimum Inoculum of Resistance Assay for Evaluating Antitoxoplasmosis Compounds That Target Phenylalanine tRNA Synthetase. ACS Infect Dis 2026, 12 (7), 2324–2331.

(31) McFadden, D. C.; Tomavo, S.; Berry, E. A.; Boothroyd, J. C. Characterization of cytochrome b from Toxoplasma gondii and Q(o) domain mutations as a mechanism of atovaquone-resistance. Mol Biochem Parasitol 2000, 108 (1), 1–12.

(32) Reynolds, M. G.; Oh, J.; Roos, D. S. In vitro generation of novel pyrimethamine resistance mutations in the Toxoplasma gondii dihydrofolate reductase. Antimicrob Agents Chemother 2001, 45 (4), 1271–1277.

(33) Hopper, A. T.; Brockman, A.; Wise, A.; Gould, J.; Barks, J.; Radke, J. B.; Sibley, L. D.; Zou, Y.; Thomas, S. Discovery of Selective Toxoplasma gondii Dihydrofolate Reductase Inhibitors for the Treatment of Toxoplasmosis. J Med Chem 2019, 62 (3), 1562–1576.

(34) Schneider-Poetsch, T.; Ju, J.; Eyler, D. E.; Dang, Y.; Bhat, S.; Merrick, W. C.; Green, R.; Shen, B.; Liu, J. O. Inhibition of eukaryotic translation elongation by cycloheximide and lactimidomycin. Nat Chem Biol 2010, 6 (3), 209–217.

(35) Kantarjian, H. M.; Chifotides, H. T.; Haddad, F. G.; Short, N. J.; Loghavi, S.; Jabbour, E. Ponatinib-review of historical development, current status, and future research. Am J Hematol 2024, 99 (8), 1576–1585.

(36) Todd, G. C.; Novilla, M. N.; Howard, L. C. Comparative toxicology of monensin sodium in laboratory animals. J Anim Sci 1984, 58 (6), 1512–1517.

(37) Ekinci, I. B.; Chlodowska, A.; Olejnik, M. Ionophore Toxicity in Animals: A Review of Clinical and Molecular Aspects. Int J Mol Sci 2023, 24 (2),

(38) Miller, S. C.; Huang, R.; Sakamuru, S.; Shukla, S. J.; Attene-Ramos, M. S.; Shinn, P.; Van Leer, D.; Leister, W.; Austin, C. P.; Xia, M. Identification of known drugs that act as inhibitors of NF-kappaB signaling and their mechanism of action. Biochem Pharmacol 2010, 79 (9), 1272–1280.

(39) Ennis, H. L.; Lubin, M. Cycloheximide: Aspects of Inhibition of Protein Synthesis in Mammalian Cells. Science 1964, 146 (3650), 1474–1476.

(40) Lane, M. E.; Yu, B.; Rice, A.; Lipson, K. E.; Liang, C.; Sun, L.; Tang, C.; McMahon, G.; Pestell, R. G.; Wadler, S. A novel cdk2-selective inhibitor, SU9516, induces apoptosis in colon carcinoma cells. Cancer Res 2001, 61 (16), 6170–6177.

(41) Cicenas, J.; Valius, M. The CDK inhibitors in cancer research and therapy. J Cancer Res Clin Oncol 2011, 137 (10), 1409–1418.

(42) Sawada, I.; Hashimoto, K.; Sawada, K.; Kinose, Y.; Nakamura, K.; Toda, A.; Nakatsuka, E.; Yoshimura, A.; Mabuchi, S.; Fujikawa, T.;, et al. The Novel IkappaB Kinase beta Inhibitor, IMD-0560, Has Potent Therapeutic Efficacy in Ovarian Cancer Xenograft Model Mice. Int J Gynecol Cancer 2016, 26 (4), 610–618.

(43) Chon, H. J.; Bae, K. J.; Lee, Y.; Kim, J. The casein kinase 2 inhibitor, CX-4945, as an anti-cancer drug in treatment of human hematological malignancies. Front Pharmacol 2015, 6, 70.

(44) Testing the Safety and Tolerability of CX-4945 in Patients With Recurrent Medulloblastoma Who May or May Not Have Surgery. 2025. https://clinicaltrials.gov/study/NCT03904862?tab=study (accessed).

(45) Gray, J. E.; Infante, J. R.; Brail, L. H.; Simon, G. R.; Cooksey, J. F.; Jones, S. F.; Farrington, D. L.; Yeo, A.; Jackson, K. A.; Chow, K. H.;, et al. A first-in-human phase I dose-escalation, pharmacokinetic, and pharmacodynamic evaluation of intravenous LY2090314, a glycogen synthase kinase 3 inhibitor, administered in combination with pemetrexed and carboplatin. Invest New Drugs 2015, 33 (6), 1187–1196.

(46) Rizzieri, D. A.; Cooley, S.; Odenike, O.; Moonan, L.; Chow, K. H.; Jackson, K.; Wang, X.; Brail, L.; Borthakur, G. An open-label phase 2 study of glycogen synthase kinase-3 inhibitor LY2090314 in patients with acute leukemia. Leuk Lymphoma 2016, 57 (8), 1800–1806.

(47) Sleda, M. A.; Diagne, K.; Clifton, V. M.; Baierna, B.; Manetsch, R.; Moreno, S. N. J. Legacy 4(1H)-Quinolone Scaffolds Activity against Acute and Chronic Toxoplasma gondii Infection. ACS Infect Dis 2026, 12 (6), 1901–1914.

(48) Cortes, J. E.; Kim, D. W.; Pinilla-Ibarz, J.; le Coutre, P.; Paquette, R.; Chuah, C.; Nicolini, F. E.; Apperley, J. F.; Khoury, H. J.; Talpaz, M.;, et al. A phase 2 trial of ponatinib in Philadelphia chromosome-positive leukemias. N Engl J Med 2013, 369 (19), 1783–1796.

(49) FDA OK’s Iclusig’s return to market. Cancer Discov 2014, 4 (2), 138.

(50) Breccia, M.; Iurlo, A.; Pane, F.; Scortechini, A. R.; Rossi, A. R.; Santoro, M.; Cerrano, M.; Lunghi, F.; Maggi, A.; Di Renzo, N.;, et al. Long-term safety and efficacy of ponatinib in intolerant chronic myeloid leukemia: subanalysis of the observational study of Iclusig(R) (ponatinib) treatment in patients with chronic myeloid leukemia in Italy. Haematologica 2026,

(51) Jabbour, E.; Apperley, J.; Cortes, J.; Rea, D.; Deininger, M.; Abruzzese, E.; Chuah, C.; DeAngelo, D. J.; Hochhaus, A.; Lipton, J. H.;, et al. Dose modification dynamics of ponatinib in patients with chronic-phase chronic myeloid leukemia (CP-CML) from the PACE and OPTIC trials. Leukemia 2024, 38 (3), 475–481.

(52) Yap, T. A.; Goldman, J. W.; Vinayak, S.; Tomova, A.; Hamilton, E.; Naito, Y.; Giordano, A.; Bondarenko, I.; Yamashita, T.; Zhou, L.;, et al. First-in-Human Phase I/IIa Study of the First-in-Class CDK2/4/6 Inhibitor PF-06873600 Alone or with Endocrine Therapy in Patients with Breast Cancer. Clin Cancer Res 2025, 31 (14), 2899–2909.

(53) Freeman-Cook, K.; Hoffman, R. L.; Miller, N.; Almaden, J.; Chionis, J.; Zhang, Q.; Eisele, K.; Liu, C.; Zhang, C.; Huser, N.;, et al. Expanding control of the tumor cell cycle with a CDK2/4/6 inhibitor. Cancer Cell 2021, 39 (10), 1404–1421 e1411.

(54) Dorne, J. L.; Fernandez-Cruz, M. L.; Bertelsen, U.; Renshaw, D. W.; Peltonen, K.; Anadon, A.; Feil, A.; Sanders, P.; Wester, P.; Fink-Gremmels, J. Risk assessment of coccidostatics during feed cross-contamination: animal and human health aspects. Toxicol Appl Pharmacol 2013, 270 (3), 196–208.

(55) Haverkos, B.; Alpdogan, O.; Baiocchi, R.; Brammer, J. E.; Feldman, T. A.; Capra, M.; Brem, E. A.; Nair, S.; Scheinberg, P.; Pereira, J.;, et al. Targeted therapy with nanatinostat and valganciclovir in recurrent EBV-positive lymphoid malignancies: a phase 1b/2 study. Blood Adv 2023, 7 (20), 6339–6350.

(56) Blain, M.; Garrard, A.; Poppenga, R.; Chen, B.; Valento, M.; Halliday Gittinger, M. Survival After Severe Rhabdomyolysis Following Monensin Ingestion. J Med Toxicol 2017, 13 (3), 259–262.

(57) Obrig, T. G.; Culp, W. J.; McKeehan, W. L.; Hardesty, B. The mechanism by which cycloheximide and related glutarimide antibiotics inhibit peptide synthesis on reticulocyte ribosomes. J Biol Chem 1971, 246 (1), 174–181.

(58) Ljubenkov, P. A.; Edwards, L.; Iaccarino, L.; La Joie, R.; Rojas, J. C.; Koestler, M.; Harris, B.; Boeve, B. F.; Borroni, B.; van Swieten, J. C.;, et al. Effect of the Histone Deacetylase Inhibitor FRM-0334 on Progranulin Levels in Patients With Progranulin Gene Haploinsufficiency: A Randomized Clinical Trial. JAMA Netw Open 2021, 4 (9), e2125584.

(59) Brown, N. R.; Korolchuk, S.; Martin, M. P.; Stanley, W. A.; Moukhametzianov, R.; Noble, M. E. M.; Endicott, J. A. CDK1 structures reveal conserved and unique features of the essential cell cycle CDK. Nat Commun 2015, 6, 6769.

(60) Tada, Y.; Kokabu, S.; Sugiyama, G.; Nakatomi, C.; Aoki, K.; Fukushima, H.; Osawa, K.; Sugamori, Y.; Ohya, K.; Okamoto, M.;, et al. The novel IkappaB kinase beta inhibitor IMD-0560 prevents bone invasion by oral squamous cell carcinoma. Oncotarget 2014, 5 (23), 12317–12330.

(61) Condie, L. W.; Buhler, D. R. Effect of hexachlorophene, tribromsalan, trichlorcarban and cloflucarban on hexobarbital sleeping time and hepatic drug-metabolizing enzyme activity in vitro in the rat. Biochem Pharmacol 1979, 28 (3), 375–380.

(62) Bota, D. A.; Mason, W.; Kesari, S.; Magge, R.; Winograd, B.; Elias, I.; Reich, S. D.; Levin, N.; Trikha, M.; Desjardins, A. Marizomib alone or in combination with bevacizumab in patients with recurrent glioblastoma: Phase I/II clinical trial data. Neurooncol Adv 2021, 3 (1), vdab142.

(63) Potts, B. C.; Albitar, M. X.; Anderson, K. C.; Baritaki, S.; Berkers, C.; Bonavida, B.; Chandra, J.; Chauhan, D.; Cusack, J. C., Jr.; Fenical, W.;, et al. Marizomib, a proteasome inhibitor for all seasons: preclinical profile and a framework for clinical trials. Curr Cancer Drug Targets 2011, 11 (3), 254–284.

(64) Di, K.; Lloyd, G. K.; Abraham, V.; MacLaren, A.; Burrows, F. J.; Desjardins, A.; Trikha, M.; Bota, D. A. Marizomib activity as a single agent in malignant gliomas: ability to cross the blood-brain barrier. Neuro Oncol 2016, 18 (6), 840–848.

(65) Kusaczuk, M.; Piskorz, W. M.; Domasik, J. Marizomib in the therapy of brain tumors-how far did we go and where do we stand? Pharmacol Rep 2025, 77 (5), 1177–1195.

(66) Wu, M.; Hui, S. Y.; Skora, A.; Chan, Y. Y.; Chung, G. T.; Li, L.; Tao, Q.; Guo, D.; Dawson, C. W.; Young, L. S.;, et al. Targeting EBV-associated gastric cancer by lytic induction therapy with nanatinostat. Tumour Virus Res 2026, 22, 200346.

(67) Akinyemi, I. A.; Zeigler, T. M.; Willman, G. H.; Patel, D.; El-Guindy, A.; McIntosh, M. T.; Bhaduri-McIntosh, S. Clinical-Stage Nanatinostat Triggers Productive Epstein-Barr Virus Lytic Replication. J Med Virol 2026, 98 (8), e71077.

(68) Nunes, D. C. O.; Bispo-da-Silva, L. B.; Napolitano, D. R.; Costa, M. S.; Figueira, M.; Rodrigues, R. S.; Rodrigues, V. M.; Yoneyama, K. A. G. In vitro additive interaction between ketoconazole and antimony against intramacrophage Leishmania (Leishmania) amazonensis amastigotes. PLoS One 2017, 12 (6), e0180530.

(69) Rosenthal, M.; Goldberg, D. E. 4-Aminoquinolines block heme iron reactivity and interfere with artemisinin action. Elife 2026, 14, RP108976.

(70) Ence, C. C.; Uddin, T.; Borrel, J.; Mittal, P.; Xie, H.; Zoller, J.; Sharma, A.; Comer, E.; Schreiber, S. L.; Melillo, B.;, et al. Bicyclic Pyrrolidine Inhibitors of Toxoplasma gondii Phenylalanine t-RNA Synthetase with Antiparasitic Potency In Vitro and Brain Exposure. ACS Infect Dis 2024, 10 (6), 2212–2221.

(71) Diaz-Martin, S.; Swale, C.; Bellini, V.; Dobrescu, I.; Wenker, J.; Brenier-Pinchart, M. P.; Braun, L.; Tollec, A.; Corrao, C.; Coute, Y.;, et al. Structural and functional characterization of TgGSK3, a druggable kinase in Toxoplasma gondii. Nat Commun 2025, 16 (1), 9765.

(72) Eldar-Finkelman, H.; Martinez, A. GSK-3 Inhibitors: Preclinical and Clinical Focus on CNS. Front Mol Neurosci 2011, 4, 32.

(73) Tobin, C. M.; Knoll, L. J. A patatin-like protein protects Toxoplasma gondii from degradation in a nitric oxide-dependent manner. Infect Immun 2012, 80 (1), 55–61.

(74) Vizcarra, E. A.; Goerner, A. L.; Ulu, A.; Hong, D. D.; Bergersen, K. V.; Talavera, M. A.; Le Roch, K.; Wilson, E. H.; White, M. W. An ex vivo model of Toxoplasma recrudescence reveals developmental plasticity of the bradyzoite stage. mBio 2023, 14 (5), e0183623.

