## Supplementary materials for "High-Throughput Screen of Advanced Repurposing Library Identifies Compounds Active Against Acute and Chronic Stages of *Toxoplasma gondii*"

**Table S1**: Sources and catalog numbers of the top 15 hit compounds.

| Compound Name | Vendor/Source | Catalog Number |
|---|---|---|
| Silmitasertib | Sigma Aldrich | A103748 |
| LY-2090314 | Sigma Aldrich | SML1438 |
| Narasin | Sigma Aldrich | N1271 |
| Monensin | Sigma Aldrich | 46468 |
| Ponatinib-HCl | Sigma Aldrich | AMBH9A90385A |
| Ebvaciclib | Aurum Pharmatech | 2185857978 |
| Cycloheximide | Sigma Aldrich | 239763 |
| EVP-0334 | Calibr collaboration | kCCW911 |
| SU-9516 | Sigma Aldrich | A137702 |
| IMD-0560 | Sigma Aldrich | TA9H9453336D |
| kCOT677 | Calibr collaboration | CBR-050-353-072-1 |
| Cloflucarban | Aurum Pharmatech | BN15713 |
| Fluorosalan | Millipore Sigma | AABH97D2BE46 |
| Marizomib | Aaron Chemicals LLC | AR00DKV2 |
| Nanatinostat | Millipore Sigma | AA01V6Y6 |

**Table S2:** $EC_{50}$ values and fold resistance of wild type (WT) and resistant populations.

| | Parasite | $EC_{50}$[a] | Fold resistance |
|---|---|---|---|
| LY-2090314 | WT | 30.90 | |
| | F1 | 408.83 | 13 |
| | F3 | 1088.81 | 35 |
| Evaciclib | WT | 60.67 | |
| | F1 | 100.00 | 2 |
| | F3 | 449.64 | 7 |
| | F3 | 54.28 | 1 |
| Nanatinostat | WT | 1.89 | |
| | F1 | 11.94 | 6 |
| | F3 | 6.98 | 4 |
| | F3 | 7.03 | 4 |

a Average vale in nM

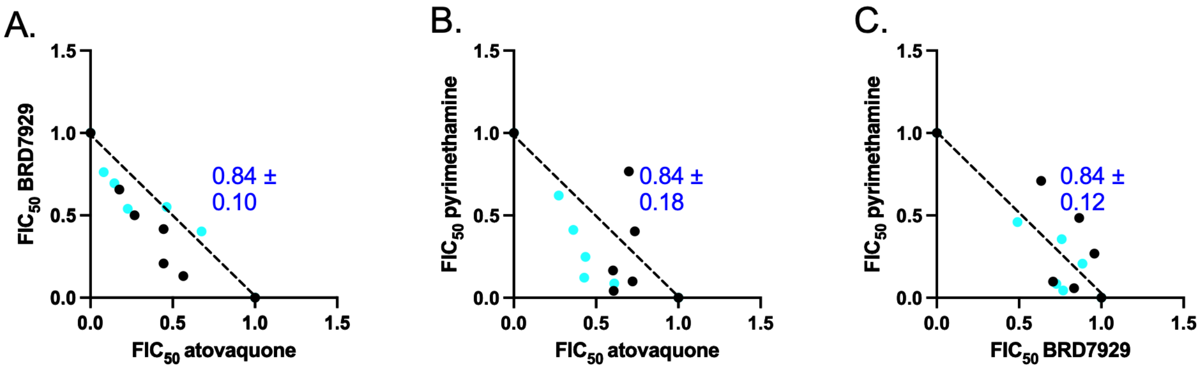


**Figure S1. Isobologram analysis of interactions among reference anti-*Toxoplasma* compounds in TgME49Fluc tachyzoites.** Fixed-ratio drug-combination assays were performed with atovaquone, BRD7929, and pyrimethamine. Drug interactions were quantified by isobologram analysis and expressed as the Fractional Inhibitory Concentration index at 50% growth inhibition ($\Sigma FIC_{50}$). Values represent the average ± SD from two independent biological replicates.
